# Subtype-Specific miRNA Expression Profiling in HPV-Associated Cervical Cancer: Insights into Oncogenic Pathways and Biomarker Potential

**DOI:** 10.1101/2025.08.13.670021

**Authors:** Nur Sabrina Abd Rashid, Ahmad Aizat Abdul Aziz, Sarina Sulong, Mohd Pazudin Ismail, Nur Asyilla Che Jalil, Marjanu Hikmah, Marahaini Musa, Daniel Roza Duski, Nazihah Mohd Yunus

**Author notes:** **Corresponding authors** (NMY).

## Abstract

Cervical cancer driven by high-risk human papillomavirus (HPV) infections, remains a global health concern. Despite advances in understanding HPV-mediated carcinogenesis, the role of microRNAs (miRNAs) in subtype-specific oncogenesis remains underexplored. This study aims to profile miRNA expression patterns in cervical cancer cell lines emphasizing subtype-specific dysregulation and its implications for diagnostic and therapeutic strategies. Four cervical cancer cell lines representing HPV16, HPV18, HPV68, and HPV-negative were profiled for miRNAs expression using NanoString nCounter™ Human V3 miRNA Panel. Differentially expressed miRNAs were identified based on fold-change ≥2.0, p-value <0.05, and false discovery rate (FDR) < 0.3. Predicted target gene of miRNAs were identified from TarBase, TargetScan, and microT-CDS. Further enrichment analysis and protein-protein interaction (PPI) network were performed using DIANA-miRPath and STRING software respectively. Subtype-specific clustering revealed 18 distinct miRNA signatures unique to different HPV subtypes with miR-205-5p and miR-125b-5p were the most up-regulated and down-regulated, respectively. Bioinformatic analyses, identified PI3K-Akt signaling as a key cancer-associated pathway implicated in tumor proliferation and survival. These findings advance understanding of miRNA-driven molecular mechanisms in HPV subtype-specific cervical carcinogenesis. miR-205-5p and miR-125b-5p are potential biomarkers and therapeutic targets toward precision medicine approaches in cervical cancer management. Further validation is warranted to integrate these findings into clinical practice.

## Introduction

Cervical cancer ranks as the fourth most common malignancy in women worldwide (1). Persistent infection with high-risk human papillomavirus (HRHPV) types is the primary etiological factor. Of the more than 200 HPV genotypes identified, approximately 14 are categorized as high-risk due to their strong association with many types of cancers (2). The progression from HPV infection to cervical cancer involves complex interactions between viral factors and host cellular mechanisms. Most HPV infections are transient and are eliminated by the immune system within two years (3,4). Persistent infection with HRHPV types, like HPV16 and HPV18, can result in viral DNA integration into the host genome. This integration disrupts normal cell cycle control and promotes malignant transformation (2).

MicroRNAs (miRNAs) have emerged as important regulatory molecules in HPV-associated carcinogenesis. These small non-coding RNA typically 22 nucleotides in length, modulate gene expression at the post-transcriptional level by binding to complementary sequences of the target mRNAs, resulting in their degradation or inhibition of translation (5). miRNAs influence a wide range of cellular processes, including proliferation, differentiation, and apoptosis (5,6). Aberrant miRNA expression has been reported in multiple cancer types, including cervical cancer, where they may function as oncogenes or tumor suppressors (7–9).

In HPV-mediated cervical carcinogenesis, integration of viral DNA, particularly the E6 and E7 oncogenes into the host genome disrupts the activity of tumor suppressor proteins p53 and pRb (10–12). This disruption promotes uncontrolled cellular proliferation and increases genomic instability. Growing evidence indicates that miRNAs participate in modulating these oncogenic pathways (13). Comparative expression profiling has revealed distinct miRNA signatures in HPV-positive cervical cancer cells relative to normal cervical epithelial cells (8,11,14).

Understanding subtype-specific miRNA expression patterns is essential, as these variations influence distinct oncogenic pathways and may guide the development of more precise therapeutic and diagnostic approaches. Current miRNA-based biomarkers are only generalized for HPV-positive cancers which limit their utility in subtype-specific applications. This study addresses that gap by using advanced profiling methods to identify deregulated miRNAs unique to individual HPV subtypes. Therefore, this article aims to explore the miRNA expression profiling in HPV-associated cervical cancer, highlighting the roles of specific miRNAs in differentiating HPV subtype and provide insight into the molecular mechanisms underlying subtype-specific carcinogenesis and their potential clinical relevance.

## Material and Methods

### Culture of cell lines

Four cervical cancer cell lines were used in this study, SiHa (HPV 16), Me180 (HPV 68), HeLa (HPV 18) and C33A (HPV-negative) (ATCC, USA). Siha, HeLa and C33A cells were cultured in Eagle’s Minimum Essential Medium (EMEM) (ATCC, USA) supplemented with 10% fetal bovine serum (FBS) (Gibco, Life Technologies), 100 U/mL penicillin and 100 µg/mL streptomycin (ATCC). Me180 cells were cultured in McCoy’s 5A Medium (ATCC, USA) with the same supplements. All cells were incubated at 37 °C in a humidified atmosphere containing 5% CO₂. Only cells at passage numbers below 10 were used for downstream analyses.

### Total RNA isolation

Total RNA was extracted using miRNeasy Mini kit (Qiagen, Germany) according to the manufacturer’s protocols. The RNA concentration and purity were determined using Multiskan SkyHigh Microplate Spectrophotometer (Thermo Fisher Scientific, USA).

### microRNA expression profiling using NanoString nCounter^TM^ Human V3 miRNA Panel

The RNA samples were subjected to the Human V3 miRNA Panel (NanoString Technologies, USA) following the manufacturer’s guidelines. The assay targeted 827 human miRNAs of clinical relevance. For each reaction, 100 ng of total RNA was used. All samples were processed in duplicate and analysed on nCounter *SPRINT*^TM^ Profiler (NanoString Technologies, USA).

### Data analysis

#### NanoString microRNA expression profiling

Raw data were processed using nSolver 4.0 software (NanoString Technologies, USA). Background correction was carried out based on the mean ± standard deviation of negative controls. Data were normalized to the geometric mean of the top 100 expressed miRNAs in each sample. Differential expression was determined using a fold change (FC) threshold of ≥ 2.0, p < 0.05, and false discovery rate (FDR) < 0.3.

#### Gene ontology and pathway enrichment

The biological significance of differentially expressed miRNAs (DEMs) was assessed using DIANA-miRPath v4.0, with pathway mapping based on the Kyoto Encyclopedia of Genes and Genomes (KEGG) database. Pathways with p < 0.05 were considered significant.

#### Target gene prediction

Potential miRNA target genes were predicted using three different bioinformatics algorithms including TarBase v8.0 (MiTG scores > 0.95), TargetScan v8.0 (cumulative weighted context++ score < −0.5) and microT-CDS (interaction score > 0.7). Genes predicted by at least two of these tools were selected for further analysis.

#### Protein-protein interaction (PPI) network analysis

Predicted target genes were analyzed using STRING (https://string-db.org/) to construct a PPI network. Cytoscape software (http://www.cytoscape.org/) was used for visualization and clustering, with MCODE applied to identify significant modules. Gene Ontology and KEGG pathway enrichment analyses for each module were performed using DAVID.

## Results

### miRNA expression profile in HPV-positive cervical cancer cell lines

Analysis identified 18 miRNAs with significant differential expression in HPV-positive cervical cancer cell lines compared with the HPV-negative control. Of these, 10 were downregulated and 8 were upregulated. The most upregulated was let-7b-5p, showing an 18.18-fold increase (p < 0.05), while miR-99a-5p exhibited the greatest downregulation with a 24.78-fold decrease (p<0.05) compared to HPV-negative cell line. Fig 1 showed the significantly deregulated miRNAs in HPV-positive cell (S1 Table).

**Fig 1.**
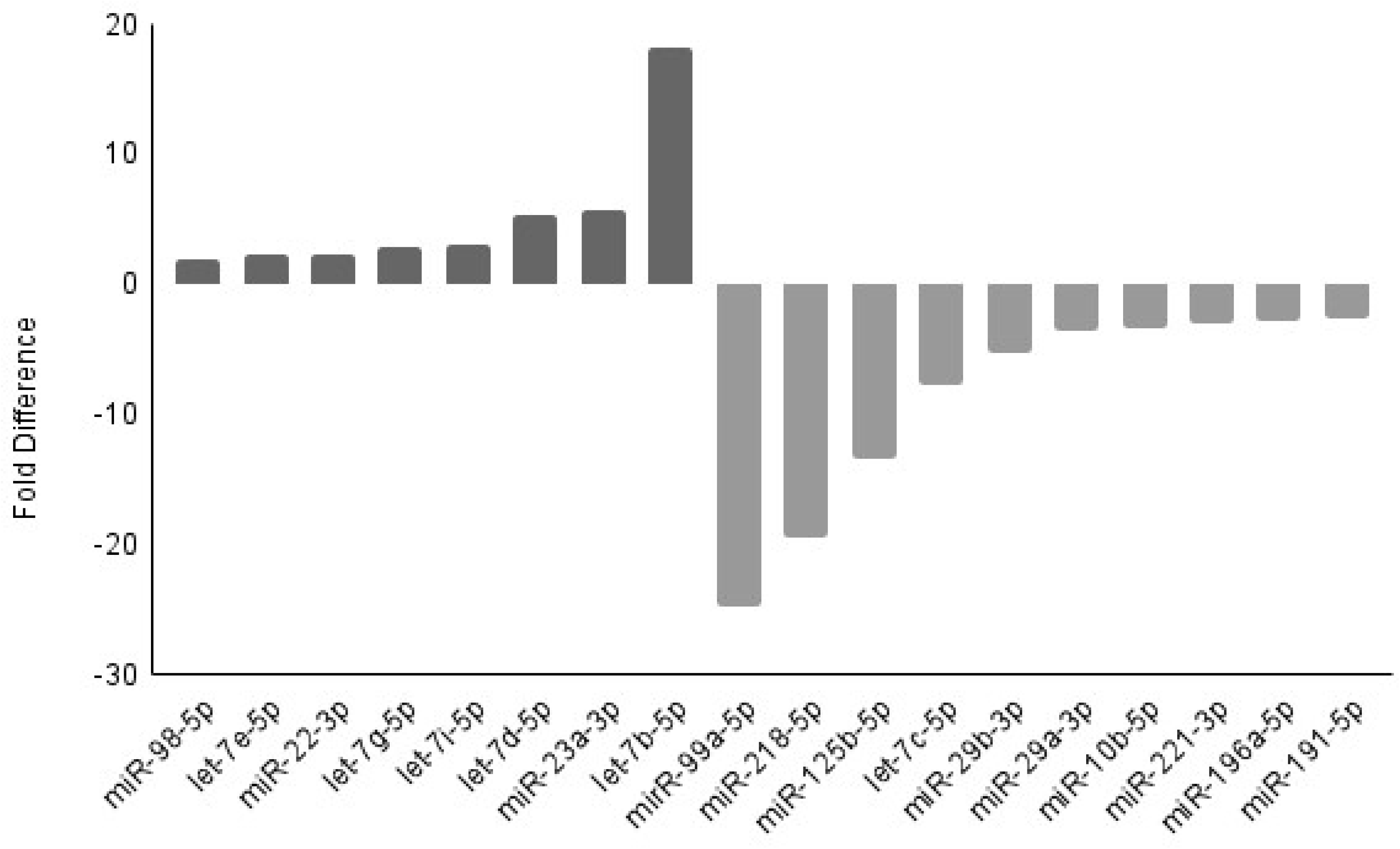
Differential expression of miRNAs in HPV-positive cervical cancer cells. The bar graph shows the fold difference in expression levels of selected miRNAs compared to HPV-negative group. Positive values indicate upregulation, while negative values indicate downregulation. Data represent the mean fold changes observed in the experimental group

### miRNAs deregulated in HR-HPV infection

An unsupervised cluster analysis between miRNA expression and HPV infection subtypes was done and showed that cell lines with different HPV subtype can be distinguished based on their miRNA expression profile (Fig 2). Number of miRNAs that are significantly altered in different HPV subtype are shown in Table 1 (S2 Table). From the results, it shows that 7 of the miRNAs are commonly expressed in all type of HPV infection. While the others 14 miRNAs showed variation in expression based on the HPV subtype as shown in Fig. 2. For HPV18, let-7f-5p (2.01-fold upregulated) and miR-221-3p (4.35-fold downregulated) were differently expressed. While only miR-181a-5p were differently upregulated by 4.66-fold in HPV16 cervical cancer cell. In HPV68 cervical cancer cell, 11 miRNAs were differently expressed where miR-205-5p and miR-125b-5p were the most upregulated (73.75-fold, p<0.05) and downregulated miRNA (44.92-fold, p<0.05) respectively.

**Fig 2.**
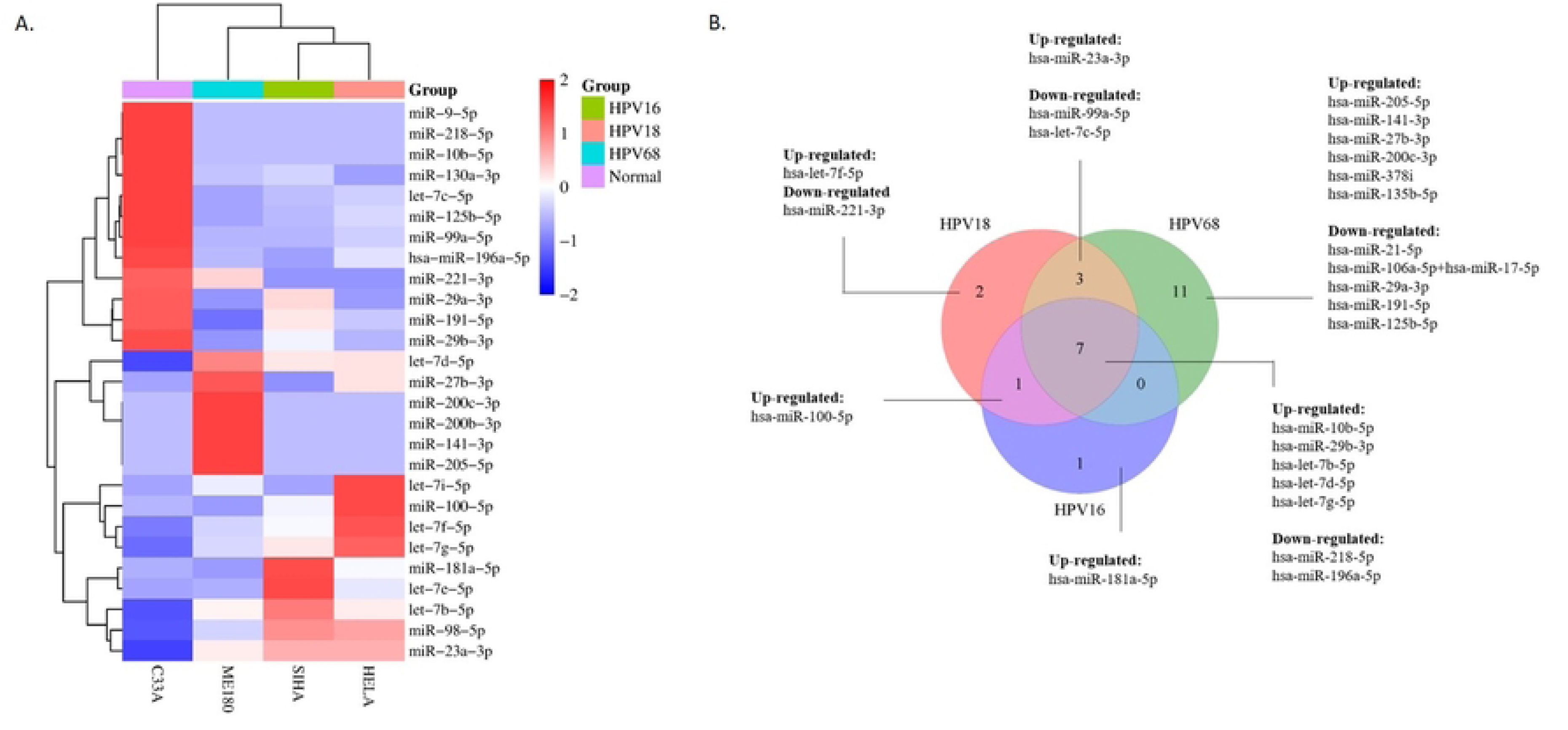
A) Hierarchical clustering heatmap of miRNA expression profiles across different cervical cancer cell lines and normal samples. Rows represent individual miRNAs, while columns correspond to cell lines: C33A (HPV-negative), ME180 (HPV68-positive), SiHa (HPV16-positive), and HeLa (HPV18-positive). The color gradient indicates normalized expression levels, with red representing upregulation and blue representing downregulation. Groups are color-coded based on HPV type, as shown in the legend. B) Venn diagram showing the overlap of dysregulated miRNAs among cervical cancer cell lines infected with HPV16, HPV18, and HPV68. Each circle represents the set of dysregulated miRNAs specific to the respective HPV type. Numbers within the overlapping regions indicate miRNAs shared between two or more groups.

**Table 1.**
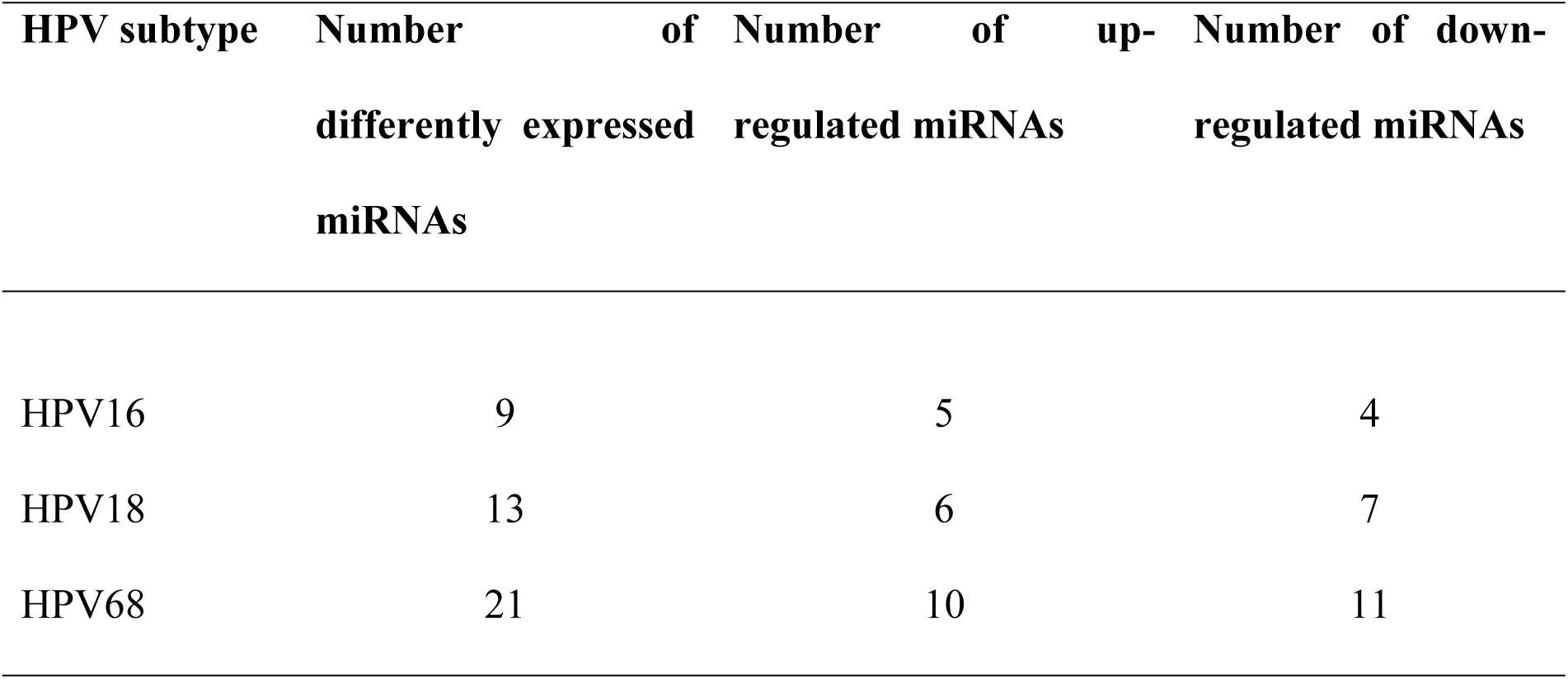
Number of significantly altered miRNAs with respect to HPV infection subtype.

### Predicted target gene regulated by the deregulated miRNAs

Target prediction using three independent bioinformatics tools identified 125 potential genes. Among the top candidates were CRKL, BCL2, CDK6, SKP2, and PTK2. The complete list of genes is provided in S3 Table.

### Gene ontology analysis of deregulated miRNAs

The gene ontology of 14 differently expressed miRNAs was analysed using DIANA-miRPath v.4. The analyses were performed separately for the up and down regulated miRNAs to see if there is any difference in their functional involvement. A *p* < 0.05 was used as a cut-off standard. The gene listed were categorised into three functional categories of gene ontology which is biological process (BP), cellular component (CC) and molecular function (MF) as shown in Fig 3 (S4 Table). There were no major differences in ontology profiles between upregulated and downregulated miRNAs.

**Fig 3.**
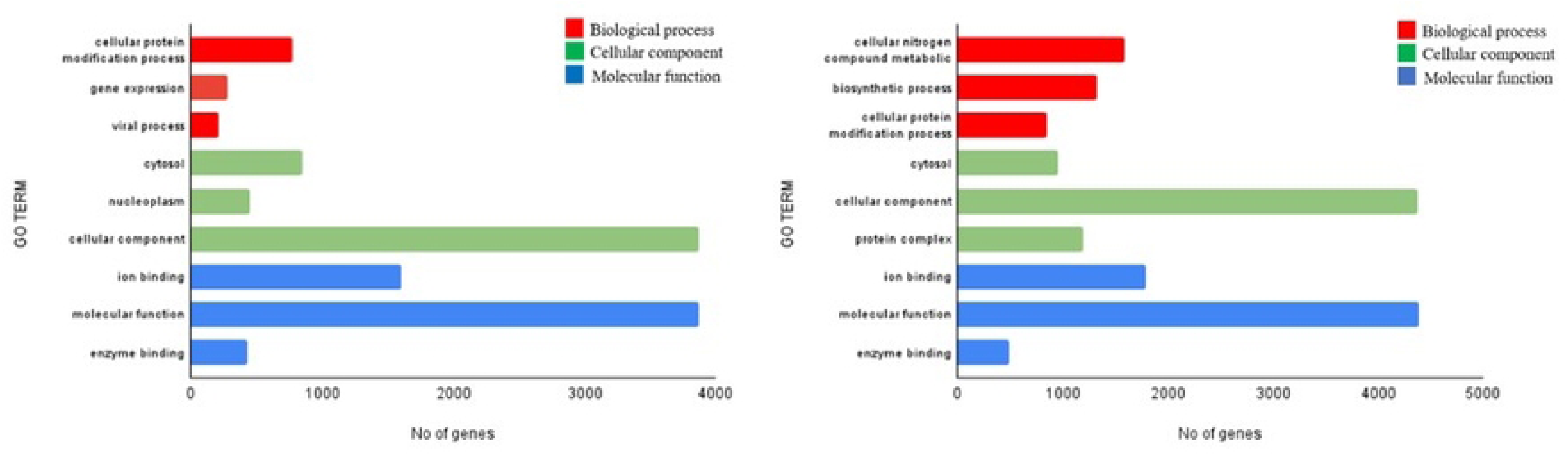
Gene Ontology (GO) analysis of the predicted target genes for miRNAs for A) Up-regulated miRNAs and B) Down-regulated miRNAs. The GO term of biological process (BP), cellular component (CC) and molecular function (MF) is distinguished by the colour code.

### Signalling pathway enrichment analysis of deregulated miRNAs

DIANA-miRPath v4.0 identified key cancer-associated signaling pathways targeted by both upregulated and downregulated miRNAs. Table 2 show top 10 important pathways targeted by the upregulated and downregulated miRNAs. It is shown that there are no different in pathways involve by upregulated and downregulated miRNAs. Both influence the important cancer-associated signalling pathways indicating a dual regulatory role of miRNAs in these processes by which this pathway showed the highest number of targeted genes in both up and downregulated miRNAs with significant p-value of 0.049 and 0.013 respectively (Fig 4, S5 Table).

**Fig 4.**
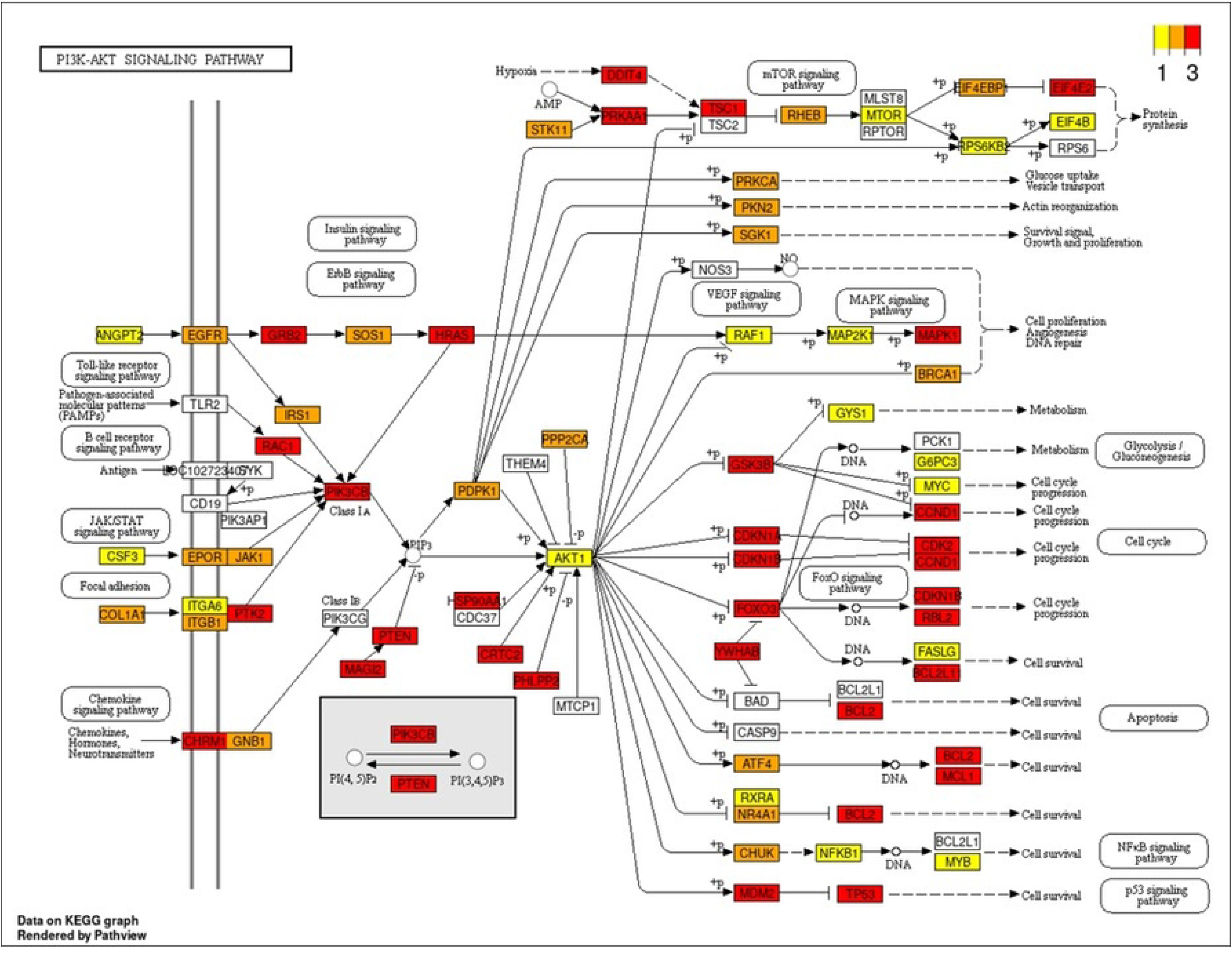
Pathway enrichment analysis of deregulated miRNAs. Figure illustrated P13K-Akt signaling pathway that shows target genes of the deregulated miRNAs such as AKT1, MTOR and BCL2. Yellow box represents target genes of one miRNAs; Orange and red box represent target genes of more than 1 miRNAs and white box indicate that the gene are not the target of these miRNAs.

**Table 2.** Top signalling pathways influenced by the signature miRNAs.

| Pathways | Name | P-value | No. Genes | No of miRNAs |
| --- | --- | --- | --- | --- |
| **Up-regulated miRNAs** |  |  |  |  |
| hsa04151 | PI3K-Akt signaling pathway | 0.049 | 115 | 7 |
| hsa04010 | MAPK signaling pathway | 0.018 | 90 | 7 |
| hsa04141 | Protein processing in endoplasmic reticulum | 7.85E-11 | 87 | 7 |
| hsa05205 | Proteoglycans in cancer | 1.79E-10 | 86 | 7 |
| hsa04144 | Endocytosis | 1.14E-05 | 85 | 7 |
| hsa04810 | Regulation of actin cytoskeleton | 0.003 | 80 | 7 |
| hsa04510 | Focal adhesion | 0.007 | 80 | 7 |
| hsa05203 | Viral carcinogenesis | 4.66E-07 | 79 | 7 |
| hsa05169 | Epstein-Barr virus infection | 0.021 | 77 | 7 |
| hsa05202 | Transcriptional misregulation in cancer | 4.53E-05 | 68 | 7 |
| **Down-regulated miRNAs** |  |  |  |  |
| hsa04151 | PI3K-Akt signaling pathway | 0.013 | 133 | 7 |
| hsa05205 | Proteoglycans in cancer | 2.44E-10 | 99 | 7 |
| hsa04010 | MAPK signaling pathway | 0.012 | 97 | 7 |
| hsa05203 | Viral carcinogenesis | 1.13E-07 | 95 | 7 |
| hsa04510 | Focal adhesion | 6.83E-05 | 95 | 7 |
| hsa04144 | Endocytosis | 0.0002 | 88 | 7 |
| hsa04141 | Protein processing in endoplasmic reticulum | 1.48E-07 | 87 | 7 |
| hsa04810 | Regulation of actin cytoskeleton | 0.006 | 86 | 7 |
| hsa05202 | Transcriptional misregulation in cancer | 0.001 | 76 | 7 |
| hsa04390 | Hippo signaling pathway | 4.38E-11 | 73 | 7 |

### Protein-Protein Interaction (PPI)network and module analysis

Of the 125 predicted target proteins, 88 formed a PPI network consisting of 553 interactions (p-value for enrichment < 1.0 × 10⁻¹⁶). Another thirty-seven proteins were excluded due to lack of network connectivity. MCODE clustering identified six significant modules within the network (Fig 5). Functional annotations for each module, including enriched GO terms and KEGG pathways, are listed in Table 3, with the complete dataset provided in S6 Table.

**Fig 5.**
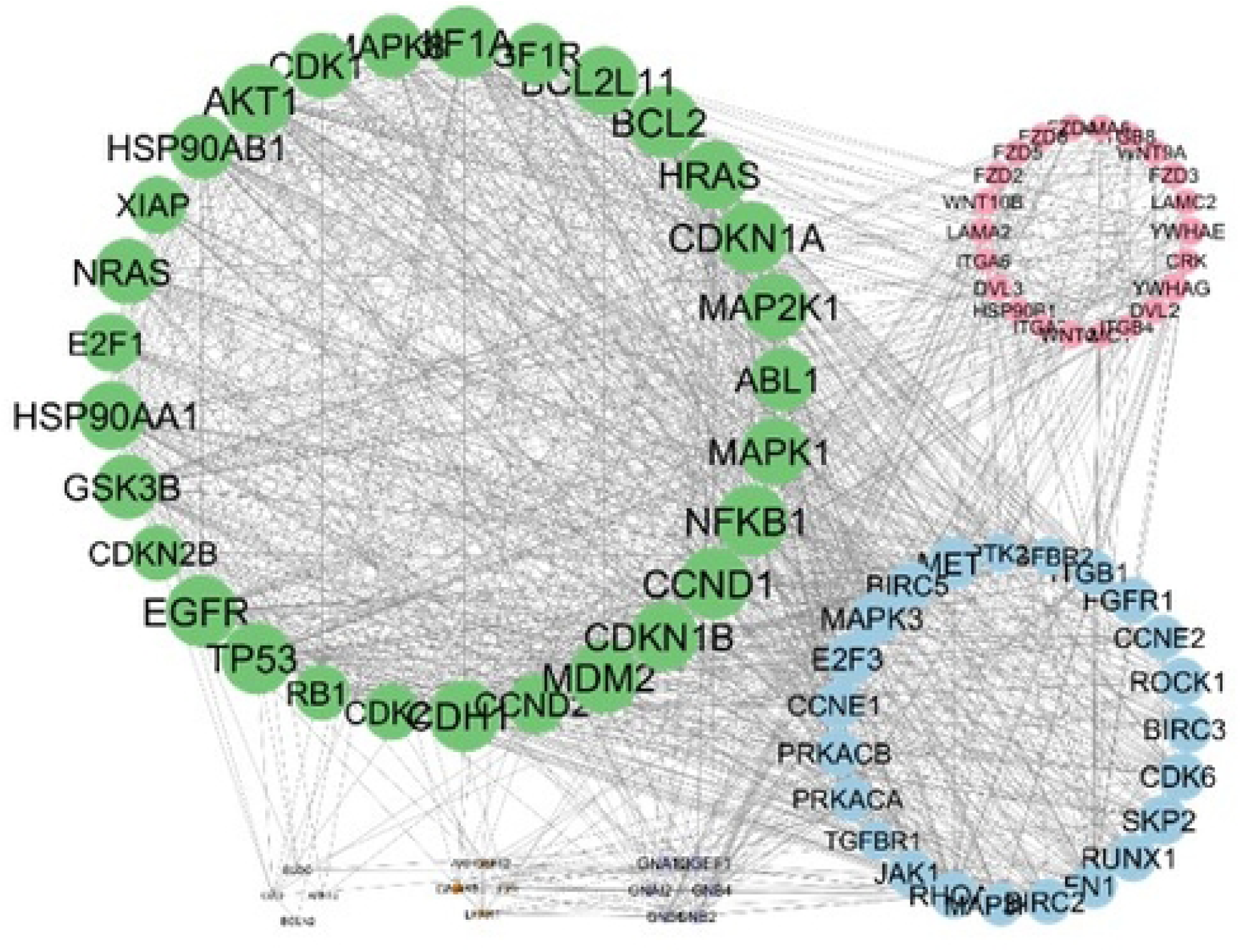
STRING protein-protein analysis of predicted target gene of deregulated miRNAs. A high number of edges indicate high intermolecular interaction between the nodes. Genes that appear bigger involved in higher and stronger interaction. Darker line shows a stronger interaction. Node fill is indicated as green (cluster 1), red (cluster 2), blue (cluster 3), purple (cluster 4), orange (cluster 5) and grey (cluster 6).

**Table 3.**
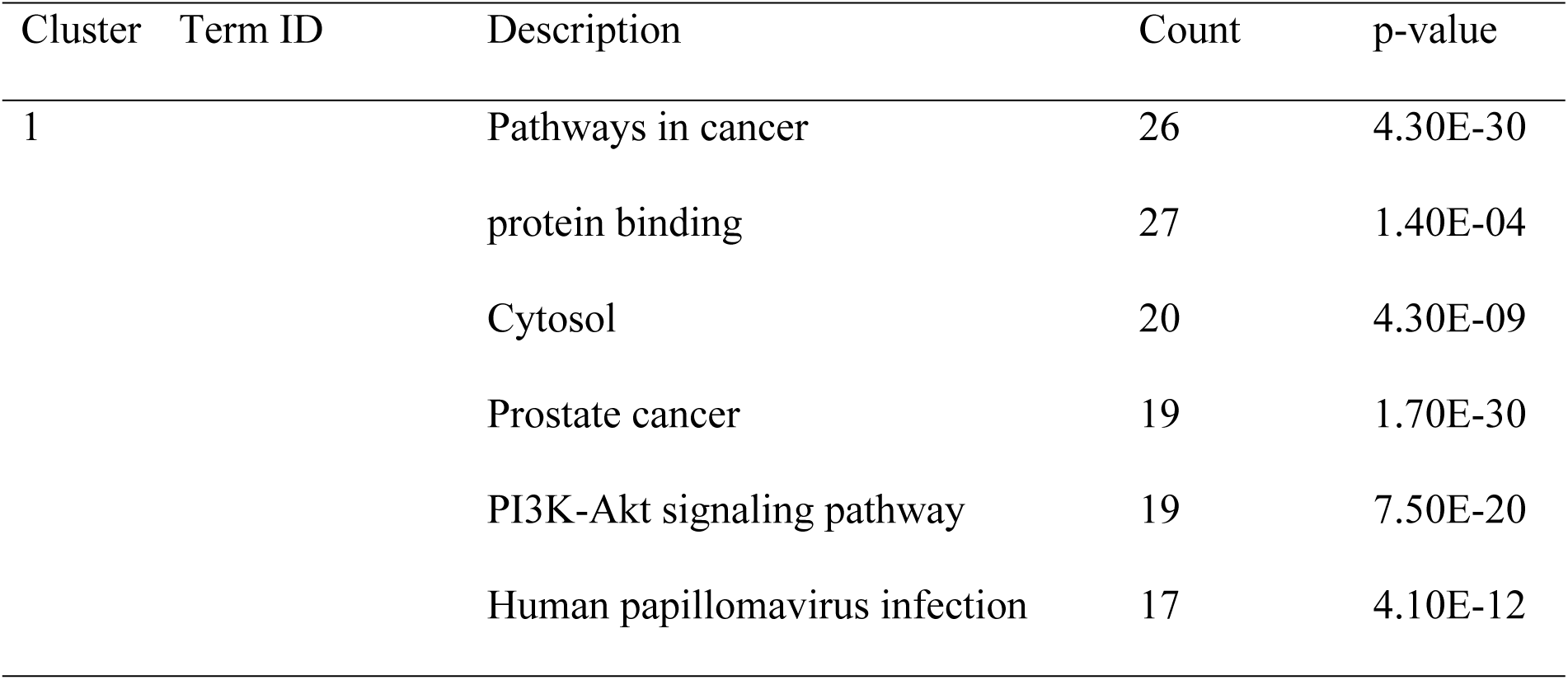
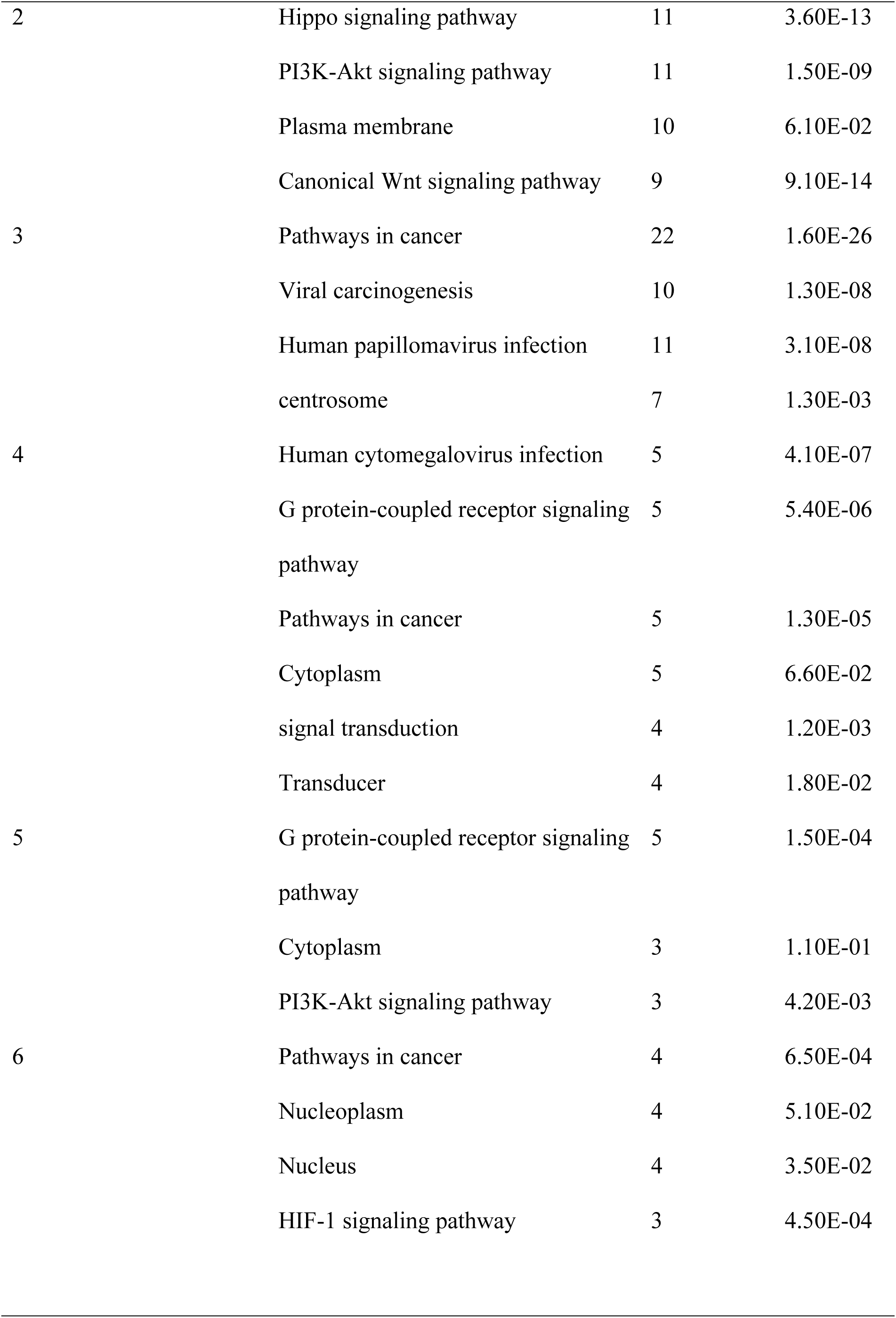
Functional annotation clustering on the clusters identified from the predicted target genes of miRNAs.

## Discussion

This study examines the influence of HPV subtypes on miRNA expression in cervical cancer, providing new insights into their role in HPV-associated disease. Expression profiling was performed in cervical cancer cell lines representing HPV16, HPV18, and HPV68 using the NanoString nCounter™ platform. Distinct miRNA expression signatures were identified across HPV-positive cells, with notable differences between subtypes (Fig 1). In addition, a subset of miRNAs was consistently deregulated regardless of HPV subtype (Fig 2). These results indicate that miRNA expression profiles are sensitive indicators of HPV infection and may also distinguish between HPV subtypes in cervical cancer.

Among the deregulated miRNAs, miR-205-5p was the most significantly upregulated, while miR-125b-5p was the most significantly downregulated. miR-205-5p showed high expression in HPV68-positive cell lines and has been linked to enhanced proliferation and migration in cervical cancer (15–17). Its overexpression correlates with advanced tumor stages and is promoted by HPV E6 and E7 oncoproteins, indicating a role in HPV-mediated oncogenesis (18). In contrast, miR-125b-5p, a known tumor suppressor, is consistently downregulated in cervical cancer tissues and HPV-positive cell lines (19,20). Its suppression, also driven by E6 and E7, likely facilitates tumor progression by removing inhibitory control over oncogenic pathways (21).

The pathway analysis revealed that several deregulated miRNAs converge on the PI3K-Akt signaling pathway, which regulates cell growth, survival, and apoptosis resistance (22,23). Dysregulation of miRNAs targeting this pathway may enable HPV-positive tumors to evade apoptosis and sustain uncontrolled proliferation. These findings align with reports that E6 and E7 oncoproteins enhance PI3K-Akt signaling to bypass p53 and Rb tumor suppressor mechanisms (24).

Several key miRNAs were implicated in this pathway, including miR-221-3p, miR-205-5p, and let-7f-5p (25,26). Downregulation of miR-221-3p in HPV18 positive cell lines suggests a tumor suppressive role, as its loss reduces PTEN expression and promotes unchecked PI3K-Akt activation (8,26–28). miR-205-5p is also known to target PTEN is significantly upregulated in cervical cancer, further sustaining PI3K-Akt signaling (16,28). It additionally enhances VEGFA expression, supporting tumor growth and angiogenesis (16,29). VEGFA upregulation strengthens PI3K-Akt activity in both cancer and endothelial cells, promoting angiogenic progression (17,30). Meanwhile, let-7f-5p targets IGF-1R, an upstream activator of PI3K-Akt, and is generally recognized as tumor-suppressor (22,23,31,32). Its increased expression in HPV-positive tumors may reflect a compensatory response to oncogenic stress which is ultimately overridden by HPV E6 and E7 activity. Together, these miRNAs regulate key nodes in the PI3K-Akt axis, contributing to HPV-driven cervical tumorigenesis (22,33).

## Conclusions

This study highlights into the differential expression profiles of miRNAs in HPV-associated cervical cancer, emphasizing the significant role of HPV subtypes in miRNA modulation. The findings reveal a subtype-specific signatures as well as miRNAs commonly deregulated across all HPV-positive cases, which not only helps in distinguishing between HPV subtypes but also underscores the miRNAs’ involvement in the PI3K-Akt signaling pathways. Clinically, identifying these subtype-specific miRNAs opens new avenues for biomarker development, enabling more precise diagnostic and prognostic tools. miRNAs such as miR-205-5p and miR-125b-5p emerged as key players, with roles in cellular proliferation, invasion, and immune evasion, thus contributing to HPV-mediated oncogenesis. While miRNAs like let-7f-5p and miR-221-3p could be used to distinguish HPV subtypes, improving early detection and subtype-specific risk stratification. Additionally, these miRNAs represent potential therapeutic targets, as modulating their expression could restore tumor suppressive functions or inhibit oncogenic pathways. Furthermore, miRNA profiling by HPV subtype could inform precision medicine approaches, tailoring treatments to address subtype-specific dysregulation, ultimately enhancing therapeutic outcomes and minimizing off-target effects. This study bridges a critical knowledge gap, providing a foundation for improved diagnostic, therapeutic, and disease-monitoring strategies in cervical cancer.

## Supporting information

**S1 Table**. List of up and down-regulated miRNAs in HPV positive cervical cancer cell lines.

**S2 Table.** List of miRNAs deregulated in HR-HPV.

**S3 Table**. List of miRNAs target gene predicted by three different algorithms.

**S4 Table**. Significantly enriched analysis of miRNAs target gene.

**S5 Table**. List of gene involved in PI3K-AKT pathway.

**S6 Table**. List of significant functional annotation clustering of PPI cluster.

